# Neuroimaging Signatures of Metacognitive Improvement in Sensorimotor Timing

**DOI:** 10.1101/2023.05.29.542773

**Authors:** Farah Bader, Martin Wiener

## Abstract

Error monitoring is an essential human ability underlying learning and metacognition. In the time domain, humans possess a remarkable ability to learn and adapt to temporal intervals, yet the neural mechanisms underlying this are not well understood. Recently, we demonstrated that humans exhibit improvements in sensorimotor time estimates when given the chance to incorporate feedback from a previous trial (Bader and Wiener 2021), suggesting that humans are metacognitively aware of their own timing errors. To test the neural basis of this metacognitive ability, human participants of both sexes underwent fMRI while they performed a visual temporal reproduction task with randomized suprasecond intervals (1-6s). Crucially, each trial was repeated following feedback, allowing a “re-do” to learn from the successes or errors in the initial trial. Behaviorally, we replicated our previous finding that subjects improve their performance on re-do trials despite the feedback being temporally uninformative (i.e. early or late). For neuroimaging, we observed a dissociation between estimating and reproducing time intervals, with the former more likely to engage regions associated with the default mode network (DMN), including the superior frontal gyri, precuneus, and posterior cingulate, whereas the latter activated regions associated traditionally with the “Timing Network” (TN), including the supplementary motor area (SMA), precentral gyrus, and right supramarginal gyrus. Notably, greater DMN involvement was observed in Re-do trials. Further, the extent of the DMN was greater on re-do trials, whereas for the TN it was more constrained. Finally, Task-based connectivity between these networks demonstrated higher inter-network correlation on initial trials, but primarily when estimating trials, whereas on re-do trials communication was higher during reproduction. Overall, these results suggest the DMN and TN work in concert to mediate subjective awareness of one’s sense of time for the purpose of improving timing performance.

**Significance Statement:** A finely tuned sense of time perception is imperative for everyday motor actions (e.g., hitting a baseball). Timing self-regulation requires correct assessment and updating duration estimates if necessary. Using a modified version of a classical task of time measurement, we explored the neural regions involved in error detection, time awareness, and learning to time. Reinforcing the role of the SMA in measuring temporal information and providing evidence of co-activation with the DMN, this study demonstrates that the brain overlays sensorimotor timing with a metacognitive awareness of its passage.

## Introduction

When learning a new task (e.g., temporal processing of interval duration) or conducting a motor movement, humans must excel at initial self-assessment and update their actions if necessary. The brain’s performance monitoring system detects errors as the motor program is performed and responds by sending a cognitive control signal to resolve the error, all without external feedback ((Ullsperger et al. 2014)). This compensatory process requires a degree of self-awareness of one’s cognitive state or metacognition ((Fleming, Huijgen, and Dolan 2012) ;(Shea et al. 2014)) Furthermore, an awareness of the subjective passage of time and the ability to self-assess one’s timing ability without prior sensory input requires error-tracking mechanisms (Kononowicz and van 2019). When extended to the sphere of learning and adapting to temporal intervals, robust systems for both error monitoring and metacognition are vital.

Past behavioral and electrophysiological evidence confirm that humans are cognitively aware of timing errors; however, questions remain concerning the extent of this time awareness ((Akdoğan and Balci 2017);(Kononowicz and van 2019); (Riemer, Kubik, and Wolbers 2019)). A recent study demonstrates that when asked to either accept or reject a trial depending on the subjective perception of proximity to the target duration interval, participants are more likely to opt out of a trial when the match (distance) between the target and reproduced time interval is lower; reduced precision in timing behavior is also observed in these trials, again supplying evidence for this self-monitoring ability ((Yallak and Balci 2022)).

Are humans aware of the direction of their timing errors (earliness or lateness) or is this cognizance limited to only error magnitude? Furthermore, how does the process of learning to time impact this awareness? Previously, we tested participants on a classical test of time reproduction, the visual time reproduction task, and incorporated feedback that did not provide information about direction (earliness or lateness) and compared it to a condition with an absence of feedback (Bader and Wiener 2021). We demonstrated that temporal estimates were more accurate and precise with post-trial non-directional feedback as participants learned and adapted to the time intervals (Bader and Wiener 2021).

Our present neuroimaging study tested the neural basis of this metacognitive ability with human participants of both sexes undergoing fMRI while they performed a visual temporal reproduction task with randomized supra-second intervals (1-6s). Crucially, each trial was repeated following feedback, allowing a “re-do” to learn from the successes or errors in the initial trial. We hypothesized that traditional time perception networks would display BOLD activation parallel to a meta-analysis of 114 human experiments performing time measurement tasks while undergoing scanning ((Cona, Wiener, and Scarpazza 2021)). The most prominent of these areas is the supplementary motor area (SMA), deemed to be the accumulator in clock time models. Other regions associated with time representation include the inferior parietal lobe (containing the supramarginal gyrus), the inferior frontal gyrus, thalamus, basal ganglia, and superior temporal gyrus ((Cona, Wiener, and Scarpazza 2021)) as well as the pre-SMA and bilateral insula (Naghibi et al. 2023).

We further hypothesized that the timing self-awareness aspect of our task may also engage the default mode network (DMN). Interactions between the timing network and default-mode-related activations are more pronounced with longer supra-second intervals (Morillon, Kell, and Giraud 2009) and could relate to mentalizing interval durations. Specifically, the posterior cingulate and the precuneus are highly implicated in self-awareness of timing ((“Neural Networks for Time Perception and Working Memory.” 2017); (Utevsky, Smith, and Huettel 2014)).

## Materials and Methods

### Participants

Twenty-seven neurologically healthy, right-handed subjects were recruited for a simultaneous fMRI-EEG experiment. EEG event markers failed to load for all stages of the task for three subjects thus leading to missing and insufficient trial counts. Another subject was unable to complete the entirety of the temporal reproduction task inside the scanner. The final data analysis included twenty-three right-handed neurologically healthy subjects (average age 23.17±4.58 SD years, 12 males, 11 females). No significant differences were observed between the ages of males (24.5±5.485) or females 21.636±2.838 according to an independent t-test (t=1.595, df=21, p=0.126)

### Task

The task was delivered via Psychopy (www.psychopy.org) from a PC desktop in the MRI console room and projected to Cambridge Research Systems BOLD Screen 32” 1920×1080 resolution (120 Hz refresh rate) screen situated ∼1.5 meters outside of the MRI Bore. The task structure of the temporal reproduction task was comprised of three phases: estimation, reproduction, and feedback for all three experiments (Figure 1). These three phases were performed twice for each duration (Bader and Wiener 2021). Each trial was initiated with a centrally presented fixation cross for a randomly presented duration of 2–6 s, drawn from a uniform distribution. In the estimation phase, a blue square was visually shown to the participant for one of five logarithmically spaced, randomly presented intervals (1.5–6 s). Until the square was on-screen, the participant was instructed to encode the duration in memory and to not use counting as a method to determine the elapsed time, which has been demonstrated as an effective means of eliminating counting strategies (Rattat and Droit-Volet 2012). Following the estimation phase, there was a 4- to 8-s gap prior to the reproduction phase, drawn from a uniform distribution. Then, the blue square reappeared on-screen in the reproduction phase and the participant was asked to press a key on an MR-compatible handheld button box (Current Designs) when the blue square had remained on-screen for the same time duration as the time elapsed in the estimation phase. The subjects’ button-press caused the square to disappear, signaling interval termination.

**Figure 1.**
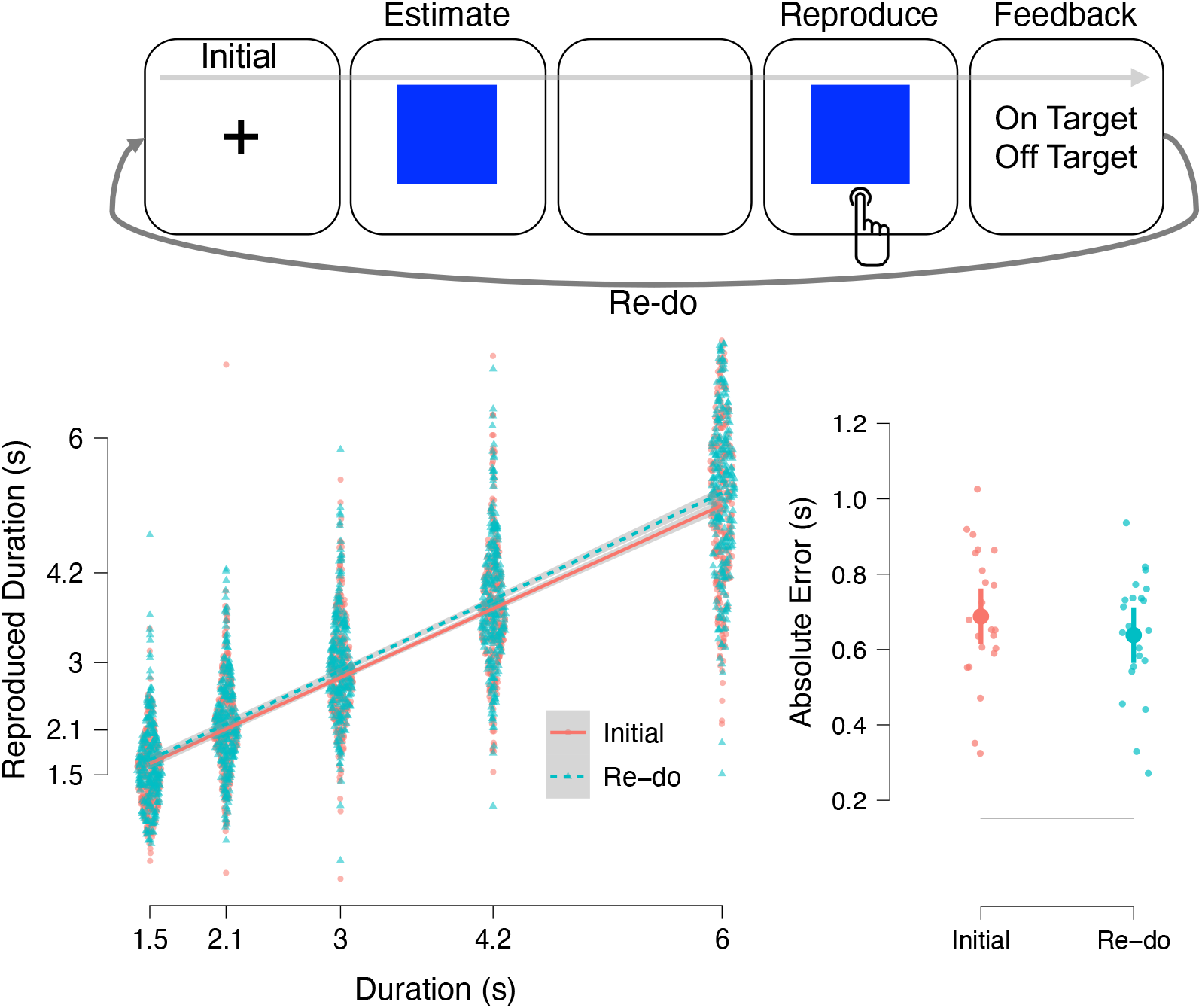
Task and data. Top: task schematic for the time reproduction task. On Initial trials, subjects were presented with a blue square for a randomly chosen interval of time, consisting of the Estimate phase. A second blue square was presented at the start of the Reproduction phase, in which subjects were required to press a response button to terminate the interval so that it matched the one just shown. Non-directional feedback was then presented, depending on whether the reproduced interval landed within an adaptive window of the presented interval. Following this, subjects were given a “Re-do” trial, on which they could repeat the entire sequence again. Bottom: Behavioral data. Left panel displays individual performance from all subjects for each of the presented durations (1-6s), in which Re-do trial performance shifted closer to the expected durations. Right panel displays the mean absolute error between Initial and Re-do trials, in which lower errors were observed on the Re-do trials; error bars represent 95% confidence intervals.

After every trial, adaptive feedback (duration = 1 second) was delivered 2–4 s after the disappearance of the square and informed the participant whether the response was on-target or off-target; notably, this feedback provided no index of temporal direction of the error (i.e. early or late). On each trial, a feedback constant (k), starting with an initial value of 3.5 was adjusted such that that the reproduced interval had to be within the window [interval/k] and was updated according to the 1-up/1-down rule with a step size of 0.015 (Jazayeri and Shadlen 2010). If the participant’s reproduced interval was either 15% above or below the target duration, on-target feedback would be delivered; otherwise, off-target feedback was delivered. Critically, after each complete trial, participants had a second opportunity (the re-do trial) to perform the entire sequence of phases (estimation, reproduction, and feedback) again, ensuring feedback was applied to the appropriate duration. After the feedback on the re-do trial, subjects experienced a random delay drawn from an exponential distribution with a minimum duration of 3s before starting the next full trial (initial and redo). In total, the experiment had 120 trials (10 durations/block × 6 blocks × two trials, initial and redo). Participants were given a break after each block for a total of six blocks.

### Yes, fMRI Acquisition

A Siemens Magnetom 3T whole-body MR scanner was used to acquire all imaging sequences. First, a localizer was performed to identify the brain’s position in space followed by a field map to measure the magnetic field inhomogeneity. Field mapping parameters were echo time (TE1)=4.92ms and TE2=7.38 ms, repetition time (TR)=731 ms, field of view (FOV)=208 mm, matrix (104×104) voxel size=2×2×2 mm and 72 slices were collected with a 2 mm thickness. Next, a structural magnetization-prepared gradient-echo-planar image (MP-RAGE) T1* was performed with the following values: repetition time (TR)=2300, echo time (TE)=2.23 ms, flip angle 8 degrees, field-of-view (FOV)=256 mm, matrix (256×256) 192 slices at a thickness of 0.88 mm were acquired with MP-RAGE sequence. Echo Planar Image (EPI) T2* scans were then collected with the following parameters (TR=2390 ms, TE=30 ms, 90-degree flip angle, FOV: 192 mm, matrix: 94×94). Forty interleaved slices with a transverse orientation, a slice thickness of 3 millimeters and 3×3×3 mm voxel size were taken. Each participant underwent one localizer, one field map, one MP-RAGE and six BOLD EPI sessions.

### fMRI Pre-Processing Steps

All pre-processing steps were performed in SPM 12 on 3D nifti files generated from the MP-RAGE, field map, and the six EPI T2 sequences. First, voxel displacement was calculated using the magnitude and phase images from the field map, followed by a slice timing correction, realignment and calculation of six affine rigid movement parameters related to head movement and unwarping. Images were then normalized to standard Montreal Neurological Institute (MNI) anatomical space and mean BOLD values were computed, written, and smoothed with a 6mm3 Gaussian kernel. Behavioral data on onset times and durations for each of the various phases (estimation, reproduction, feedback) in both the initial and redo trials were extracted and loaded by an in-house batch processing Matlab script which then estimated and wrote a first order General Linear Model (GLM) for each individual participant using the BOLD images; individual events were time-locked to the start of estimation and reproduction phases for initial and re-do trials, separately, by convolving the canonical hemodynamic response function with a boxcar stretched to match the duration of either the presented or reproduced interval on each trial (Mumford et al. 2023). Next, SPM’s contrast manager was used to create contrasts to examine changes in BOLD activation between the task phases and between the initial and redo trials of those phases in this combined dataset. The specified contrasts and weights were then fed into an additional in-house Matlab script and a second-level group GLM was estimated and written. Afterwards in SPM, one-sample t-tests were performed for each contrast and whole brain group level results were displayed with a voxelwise threshold of p<0.001, uncorrected and a cluster threshold of p<0.05, familywise error corrected.

### Statistics

JASP 0.16 (JASP team, 2021) was used to analyze the behavioral data from the temporal reproduction task. The data was normally distributed and passed the Shapiro–Wilks test of normality for the absolute temporal error, accuracy, and precision (as measured by the coefficient of variation (CV). The absolute temporal error was calculated as the reproduced duration minus the target duration. The CVs were calculated as the standard deviation of the mean reproduced durations divided by the participant’s mean reproduced durations and two separate CV values were generated for the initial and redo trials. A linear mixed effects model (fixed effects: duration, trial designation, target duration; random effects: subjects) was used to detect changes in absolute temporal error between initial and redo trials. Outlier values that were three scaled standard absolute median deviations from the median were removed.

### Functional Connectivity

To further explore putative interactions between the DMN and regions involved in time perception, we conducted an additional functional connectivity analysis. To accomplish this, we began by taking a beta-series approach, wherein single-subject GLMs were reconstructed for each subject as described above, with the exception that each trial was modeled as a separate covariate. The resulting GLM thus generated a time-series of beta values for each trial type (initial, re-do) and phase (estimation, reproduction). Next, beta-series were averaged across a series of regions of interest (ROIs). For the DMN, these regions were derived from the Yeo atlas, accessible at https://surfer.nmr.mgh.harvard.edu/fswiki/CorticalParcellation_Yeo2011 and consisted of 12 regions including the bilateral posterior cingulate cortex, superior frontal gyrus, angular gyrus, precuneus, anterior cingulate cortex, and middle temporal gyrus. All ROIs were constructed from the automatic anatomic labeling (AAL) atlas definitions. For time perception, we relied on a series of regions derived from several neuroimaging meta-analyses and recently synthesized into a series of 17 ROIs, accessible at https://neurovault.org/collections/13081/. These regions, now referred to as the Timing Network (TN), include the bilateral frontal operculum, SMA, insula, inferior parietal lobe, caudate, putamen, cerebellum (dentate gyrus), and thalamus, as well as the left precentral gyrus, right pars triangularis, and supramarginal gyrus.

To explore connectivity between these regions and how they might change across task conditions, we first calculated Spearman correlation matrices for the beta-series from all 29 ROIs, across the four task conditions; this analysis thus yielded both inter- and intra-network connectivity measures between DMN and TN regions. The Spearman correlation coefficient was chosen to reduce possible confounds driven by possible outlier trials within the beta-series. Next, to compare correlation matrices, we converted Spearman correlations to z-scores using Fisher’s r-to-z tranform. Lastly, paired t-tests were conducted comparing z-scores from each of the conditions in a 2×2 design. To assess significance, p-values were further corrected for multiple comparisons using the false-discovery rate (FDR) algorithm to a corrected value of p<0.05.

## Results

A linear mixed effects model with trial-type (Initial, Re-do) as a fixed effect and subject as a random effect, performed on the absolute temporal error, defined as the difference between reproduced and actual time interval, exhibited a significant effect of trial-type, [*F* (1,2720.04)=5.016, *p*=0.025; Model terms nested with Satterthwaite method]. Fixed effects estimates demonstrated a reduction in error on Re-do trials compared to Initial ones [*β*=-0.25, SE=0.011] such that timing performance improved when provided a second chance (Figure 1).

A repeated measures ANOVA on mean reproduced intervals revealed that the participants’ reproduced durations for the Re-do trials were also more accurate, thus nearer to the target duration and the identity line when compared to the initial trial, [F (1,22) =14.146, p<0.001] again showing an improvement in temporal estimates (Figure 1). Participants, however, did not display significantly better precision (lower coefficient of variation) in their redo trials [F (1,1.562), p<0.225, p<0.225], but did exhibit a main effect of duration F (4,26.239) p<0.001.

### Imaging Results

### Estimation Initial – Reproduction Initial Contrast

BOLD activations were seen in the bilateral superior frontal gyrus, right middle frontal gyrus, left angular and supramarginal gyrus (SMG), left middle occipital gyrus, right post- and pre-central gyrus, bilateral precuneus and posterior cingulate gyrus. In this inter-phase contrast, more parietal involvement for time perception (angular gyrus and SMG) along with pre-central gyrus for motor movement was observed in conjunction with the recruitment of structures associated with the default mode network (posterior cingulate gyrus) and metacognition (precuneus).

#### Reproduction Initial – Estimation Initial Contrast

Significant activations were observed in the left pre- and postcentral gyrus; bilateral SMA; bilateral middle cingulate gyrus; bilateral superior frontal gyrus, left central operculum, left parietal operculum, left supra-marginal gyrus, right occipital pole, right cuneus, right calcarine and lingual cortex, and superior occipital gyrus. The SMA is recruited again when re-creating the interval duration jointly with timing-related, parietal brain areas to include the SMG, left central, and parietal operculum. High detection of activity in the superior frontal gyrus also reiterates the brain’s performance monitoring system is online. Visual processing of the stimuli is emphasized again due to the eliciting of the BOLD signal in the occipital lobe.

**Figure 2.**
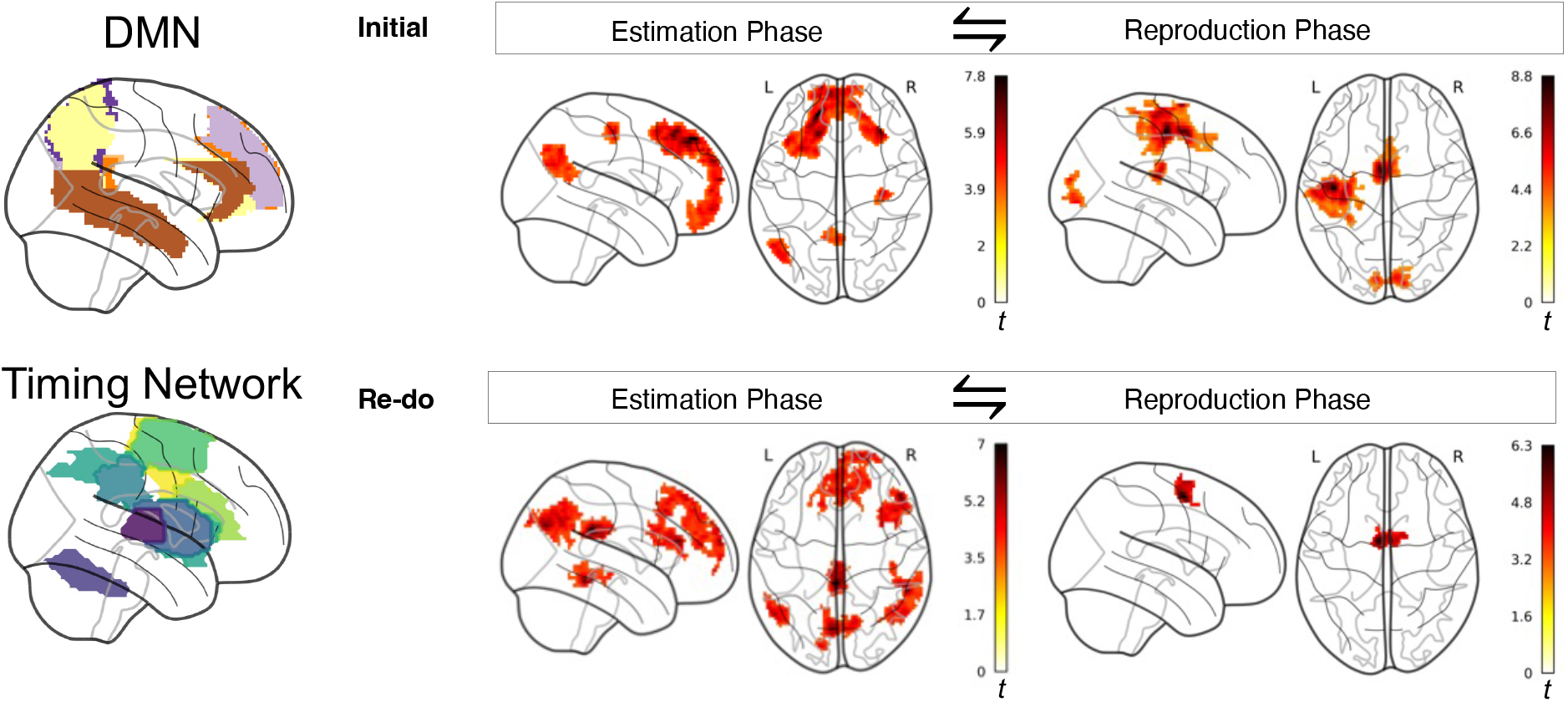
Neuroimaging results. At left: example ROIs for the default mode network (DMN) and the Timing Network are displayed. At right: the top panel displays the contrast between Estimation and Reproduction phases on Initial trials. Here, the Estimation phase invoked greater activity in DMN regions, including the superior frontal gyri, precuneus, and ACC. For the reproduction phase, activation was instead observed in the Timing Network, including the SMA and left precentral gyrus, as well as the occipital cortex. The top panel displays the same contrast, but for the Re-do trials. DMN activation was again observed for the Estimation phase, but more widespread, whereas Timing Network activation was more constrained, with only the SMA exhibiting significant activation. All displayed maps were thresholded at p<0.001, uncorrected voxelwise, and p<0.05 FWE corrected at the cluster level.

**Figure 3.**
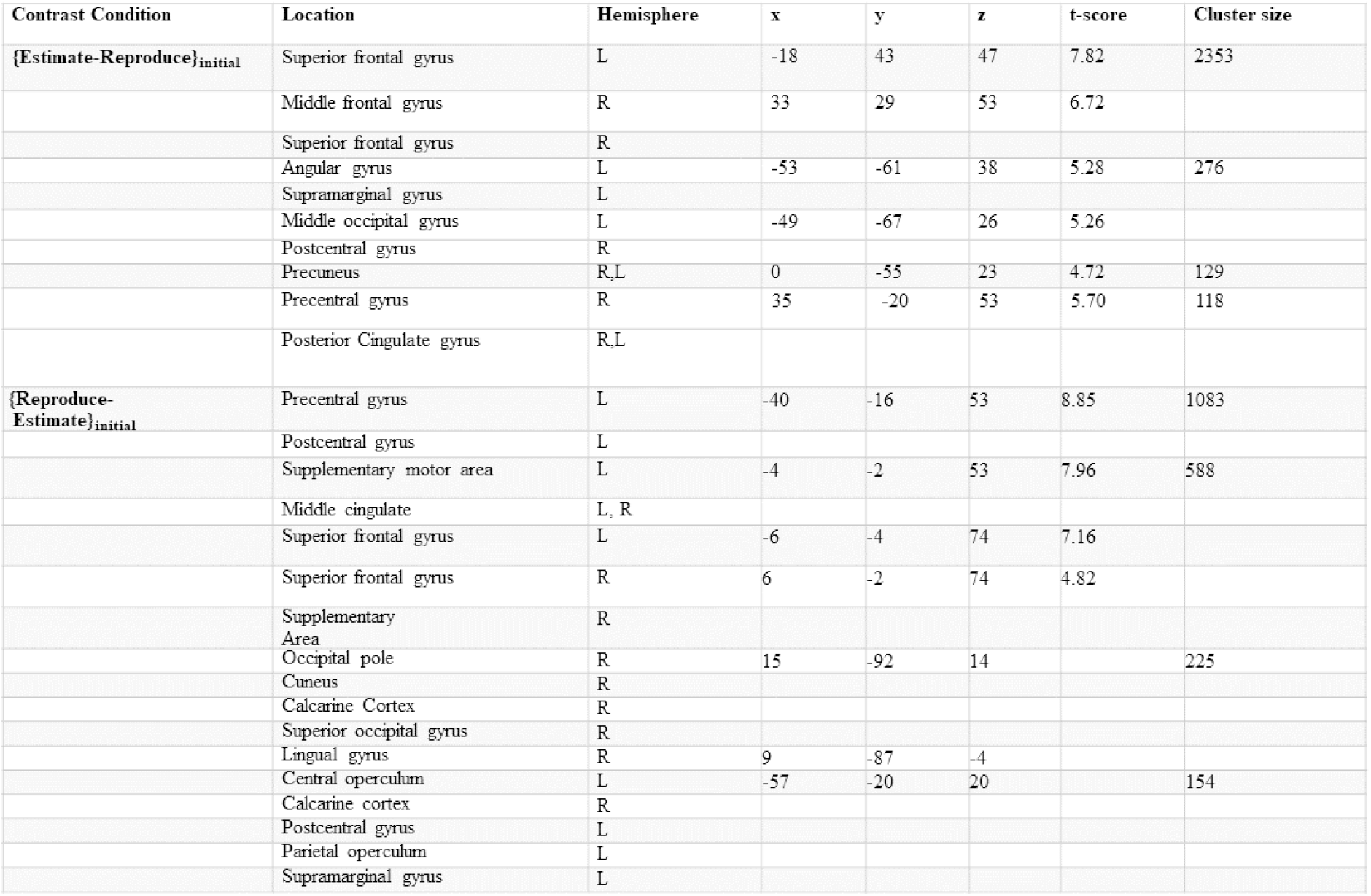
Table 1 - Estimate - Reproduce for Initial trials. MNI Coordinates are provided for all cluster peaks and sub-peaks. Brain regions lacking coordinates represent single-peak clusters overlapping multiple areas.

#### Estimation Redo - Reproduction Redo Contrast

Task related activations were witnessed in the right frontal pole, bilateral superior frontal gyrus medial segment, anterior cingulate cortex, right superior frontal gyrus, right middle frontal gyrus, right triangular and opercular part of the inferior frontal gyrus, right precentral gyrus, left superior occipital gyrus, right inferior temporal gyrus, right middle and superior temporal gyri, left cuneus and bilateral middle cingulate gyrus, bilateral precuneus and posterior cingulate respectively, bilateral angular gyrus, and left supramarginal gyrus. Regions related to forming duration judgements (inferior frontal gyrus, bilateral angular gyrus and SMG) along with self-awareness (precuneus) and the default mode network (posterior cingulate cortex) have high BOLD activation when comparing the redo trials between phases.

#### Reproduction Redo – Estimation Redo Contrast

BOLD activations were seen in the bilateral SMA and the right superior frontal gyrus, in conjunction with invoking the brain’s performance monitoring system (superior frontal gyrus).

#### Estimate Initial – Estimate Redo Contrast

No suprathreshold clusters in the fMRI peak activations for the Estimate Initial-Estimate Redo contrast were observed.

#### Estimate Redo-Estimate Initial Contrast

The fMRI Peak activations for the contrast Estimate Redo – Estimate Initial displayed high BOLD activations in the bilateral calcarine cortex, bilateral lingual gyrus, left cuneus, left occipital pole, and the left occipital fusiform gyrus. These are all visual processing areas and indicate that the subject is observing and fixating on the blue square in order to encode it.

#### Reproduce Initial – Reproduce Redo

Neural activations in the Reproduce Initial – Reproduce Redo contrasts were detected in the calcarine cortex, the exterior cerebellum, bilateral SMA (mainly right hemisphere), right superior frontal gyrus, the right anterior cingulate gyrus, left postcentral gyrus, left supramarginal gyrus, right calcarine cortex, bilateral thalamus, posterior insula, bilateral caudate, right putamen, right hippocampus, right posterior insula, right pallidum, right accumbens. Here, in addition to time-perception related areas (SMA, supramarginal gyrus, basal ganglia), and interoceptive awareness (insula), activation was observed in memory-related regions (hippocampus) as the encoded time is recalled and performance is monitored (superior frontal gyrus and anterior cingulate cortex) when decisions about the duration are made in the initial and redo trials. Notably, activation was also exhibited in the cerebellum, a region that has been associated with error learning. Previously, higher cerebellar activity has been viewed in fMRI studies in trials with sensory errors than trials without errors ((“Neural Correlates of Reach Errors” 2005); (Schlerf, Ivry, and Diedrichsen 2012)). The cerebellum also plays a role in performance monitoring, error processing, and feedback learning ((Peterburs and Desmond 2016)) in conjunction with predictive timing and the regulation of trial-by-trial variation in self-timing ((Tanaka et al. 2020)), particularly due to connections with the frontoparietal cortices ((Tanaka et al. 2020)).

#### Reproduce Redo-Reproduce Initial Contrast

No significant activation was observed using our threshold. However, using a lower threshold of p<0.001, uncorrected with a cluster extend of k=10, found task-related BOLD activations in the bilateral medial frontal cortex, bilateral gyrus rectus, bilateral superior frontal gyrus medial segment and the left medial orbital gyrus. Recruitment of brain regions in performance monitoring (medial frontal cortex) and working memory (superior frontal gyrus) ((Alagapan et al. 2019)) are displayed in the reproduction phase, demonstrating a mechanism for error detection and correction.

#### Functional Connectivity

The functional connectivity analysis revealed a double-dissociation between task phase (Estimation, Reproduction) and trial type (Initial, Re-do). Broadly, we observed that connectivity between the DMN and TN was greater in the Estimation phase, but more so on Initial trials than Re-do, whereas in the Reproduction phase, inter-network connectivity was higher on Re-do trials than Initial ones (Figure 4). More specifically, in the Estimation phase, connectivity increased between the bilateral superior frontal gyrus and ACC of the DMN and the cerebellum, left caudate and insula, inferior frontal gyrus, and left precentral gyrus of the TN; notably, the right cerebellum exhibited the largest connectivity with the DMN across a variety of structures including the angular gyrus and precuneus. In contrast, in the Reproduction phase, connectivity increased between the precuneus and the right caudate, as well as the right ACC and a variety of TN structures including the SMA, right supramarginal gyrus, and putamen. Finally, connectivity again increased between the SFG and the cerebellum, caudate, and inferior frontal gyrus.

**Figure 4.**
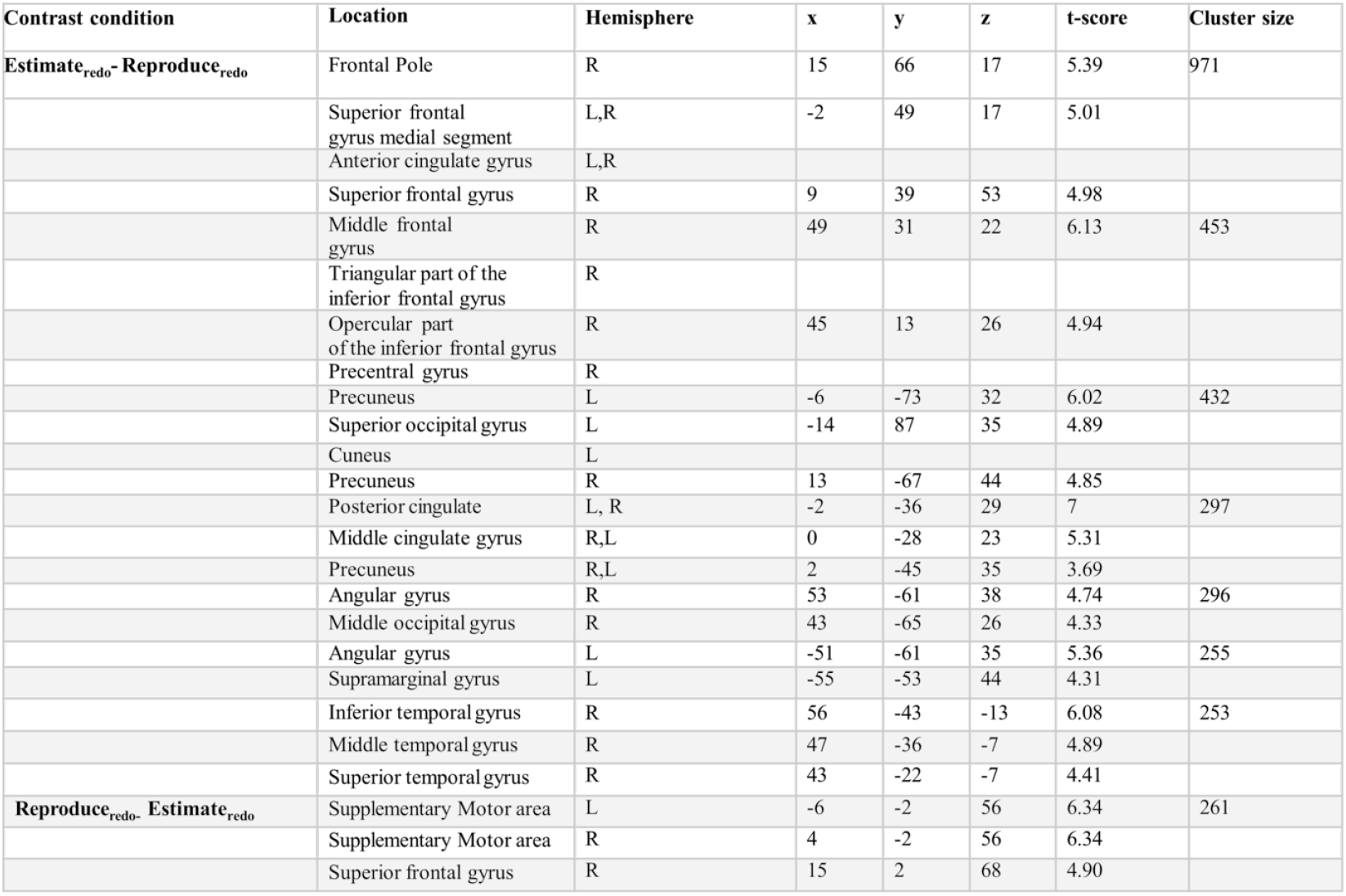
Table 2 - Estimate - Reproduce for Re-do trials. MNI Coordinates are provided for all cluster peaks and sub-peaks. Brain regions lacking coordinates represent single-peak clusters overlapping multiple areas.

**Figure 5.**
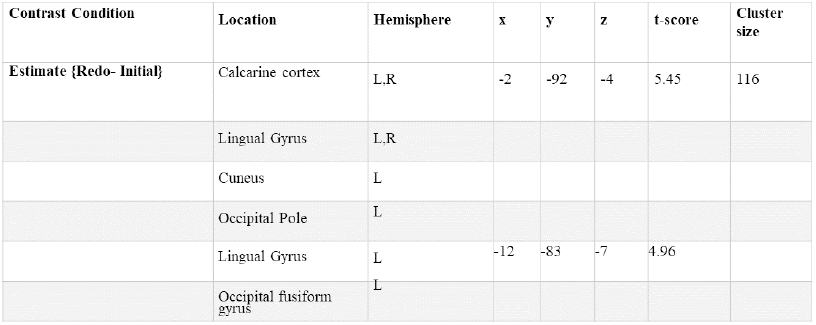
Table 3- Estimate Redo-Estimate Initial. MNI Coordinates are provided for all cluster peaks and sub-peaks. Brain regions lacking coordinates represent single-peak clusters overlapping multiple areas.

**Figure 6.**
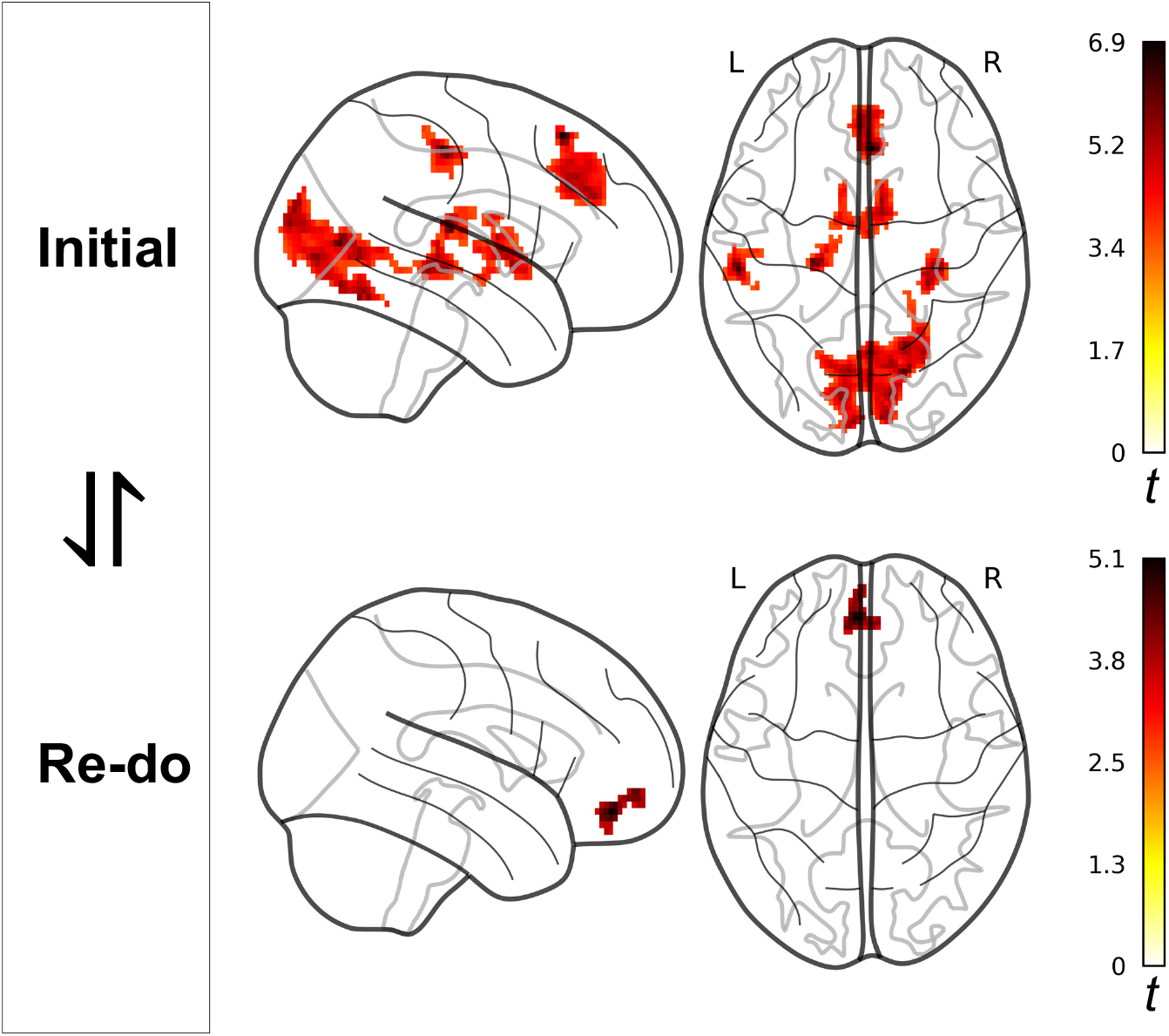
Comparisons between Initial and Re-do trials. Here, significant differences were only observed during the reproduction phase, in which the SMA, ACC, basal ganglia, and right hippocampus were active, whereas on Re-do trials only the medial orbitofrontal cortex was active.

**Figure 7.**
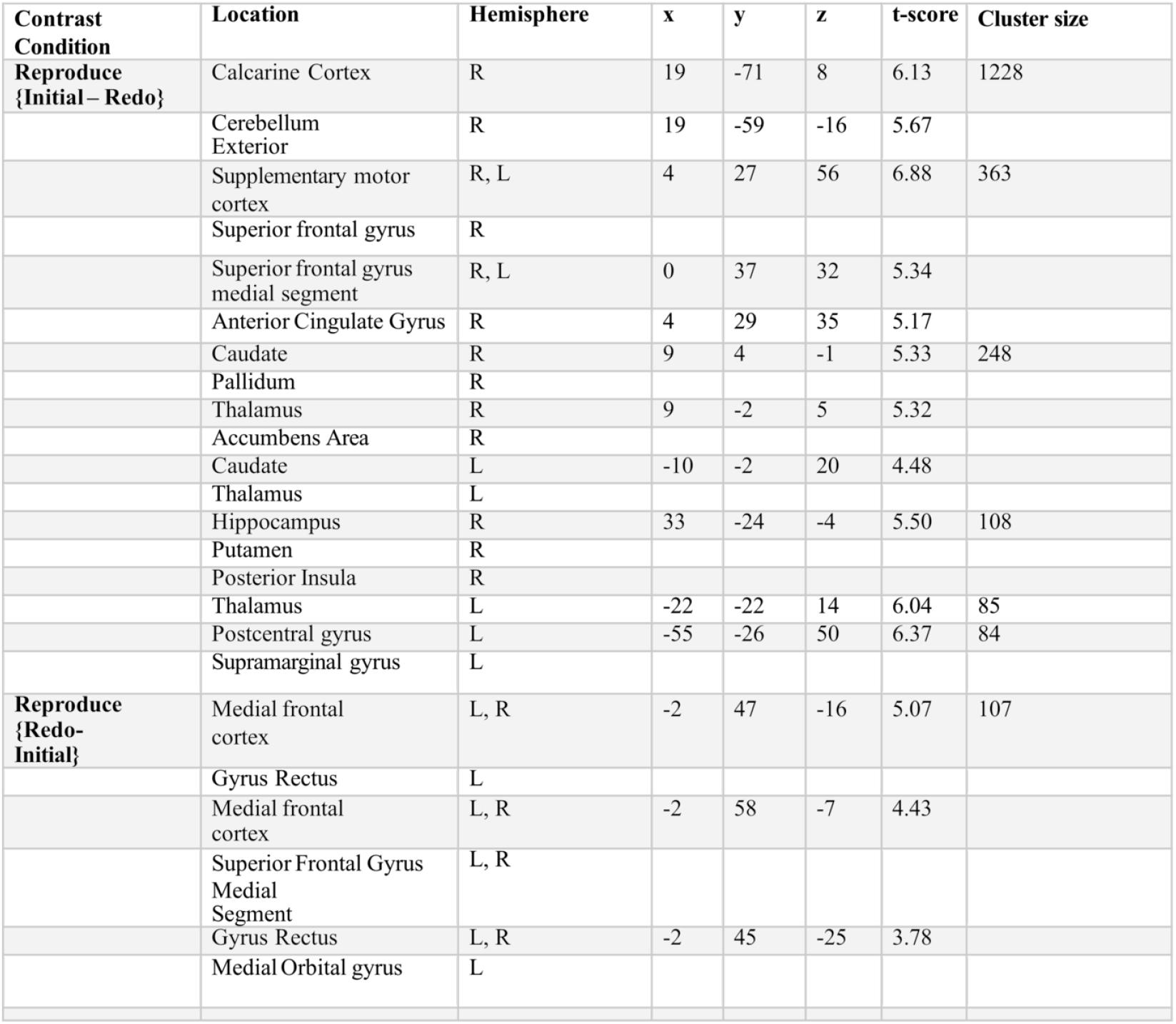
Table 4 - Initial - Re-do for Reproduction phase. MNI Coordinates are provided for all cluster peaks and sub-peaks. Brain regions lacking coordinates represent single-peak clusters overlapping multiple areas.

**Figure 8.**
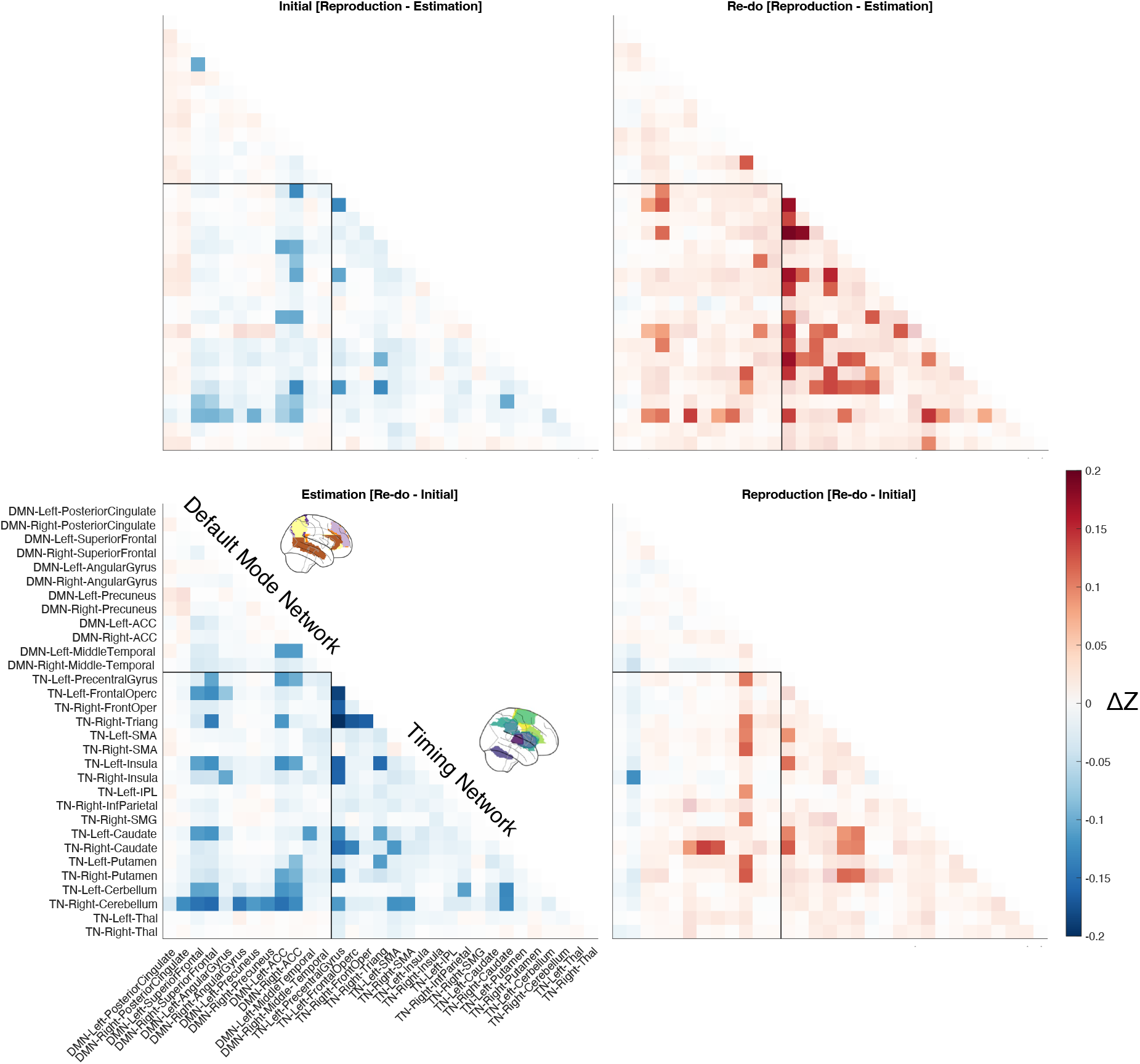
Functional connectivity results. A double-dissociation was observed across phases (Estimation vs Reproduction) and trials (Initial vs Re-do) for intra-network connectivity between the DMN and Timing Network, demarcated by the black rectangle in each panel. Here, greater connectivity was observed between the DMN and Timing Network during the Estimation phase, but primarily on Initial trials, rather than Re-do. In contrast, during the Reproduction phase, greater inter-network connectivity was observed primarily on Re-do trials. Pixel values represent the mean difference score between conditions; non-faded pixels indicate significant differences corrected for multiple comparisons with FDR to p<0.05).

## Discussion

Behavioral and electrophysiological evidence suggest that humans and rodents can monitor errors during timing behavior ((Akdoğreproduction task when tested on or off dopaminergic medicationan and Balci 2017) ; (Kononowicz and van 2019); (Kononowicz, van, and Doyère 2022)) 2022). Studies reveal that subjects are cognitively aware of the timing errors they make, particularly when learning to time interval durations, which happens rapidly and within one trial (Simen et al. 2011). Our previous behavioral study demonstrated that when allowed to “re-do” a trial, humans can incorporate non-directional feedback to improve timing estimates both in accuracy and in precision (Bader and Wiener 2021). Novel in using a mixed range of interval durations rather than a single duration, this experiment also showed that in the absence of feedback, the accuracy of time reproductions improved whereas the precision did not. These results revealed that humans are aware of the direction of their timing error, a capacity requiring metacognition; however, feedback was needed to also make timing more precise. Our current neuroimaging experiment was used to determine the neural regions underlying this improvement in timing estimation. Behaviorally, we replicated our previous finding, the absolute temporal error decreased, and the accuracy of the temporal estimates was improved with reproduced targets approaching their target durations.

Our imaging results reinforced the behavioral results but also reiterated the importance of the SMA in representing temporal information and measuring time, which has been demonstrated repeatedly in the time perception literature (Ferrandez et al., 2003;Coull, 2004; Pouthas et al., 2005;Macar et al., 2006). The SMA contains a chronotopic map with localized regions preferentially responding to specific durations along an antero-posterior gradient and exhibits duration tuning ((Protopapa et al. 2019)). As the duration is encoded during estimation, the SMA is initially recruited and invoked again in the Redo trials during reproduction. This finding is aligned with both the behavioral data and the sharpening of temporal responses as evidenced by the reduction of temporal errors and accuracy improvement in the second Redo trial.

A second focus of our study was on regions of the DMN. While studies of time perception rarely implicate the DMN, some work has suggested a link between the two. For instance, DMN activation has been shown to increase as subjects time progressively longer intervals (Morillon, Kell, and Giraud 2009), and periodic motion expectation also engages the DMN ((Carvalho et al. 2016)). More recently, multivariate lesion-symptom mapping has linked impairments in time awareness (i.e., the awareness of time passing) to disruption in DMN nodes (Skye et al. 2022). These findings suggest that the DMN and Timing Network are linked and informative to one another in the support of awareness and understanding of our sense of time. As a final note, the DMN and Timing Network largely do not overlap (Menon 2023).

Notably, our results demonstrated a dissociation between activation of the DMN and regions putatively involved in timing across the initial and Re-do trial opportunities, as well as between task phases. Specifically, estimating a duration was more likely to invoke DMN regions, whereas reproducing a duration was more likely to invoke timing regions. This difference suggests that, when explicitly encoding a time interval into memory, subjects are actively aware of time’s passage, yet when reproducing that interval the motor system drives the transformation of that estimate into a timed action. In the context of improvements in performance, DMN regions were more widespread observed on the Re-do trials than the Initial ones; yet, timing network regions were more constrained on Re-do trials. Indeed, only the SMA was significantly active on the the reproduction phase of Re-do trials. Yet, when comparing activity directly between Initial and Re-do trials, the reproduction phase also invoked activity in the hippocampus initially and the medial orbitofrontal cortex on Re-do trials. Altogether, the results suggest that DMN regions may serve to monitor timing performance and, when necessary, provide an update to motor system regions for the purpose of executing a precisely timed response.

One exception to the above findings is activation observed in the visual cortex and hippocampus during the reproduction phase. Both regions represent those outside commonly reported areas in explicit time perception, yet have both been reported to be important. For the visual cortex, previous work has shown linear increases in BOLD activation in this region in anticipation for an upcoming target (Bueti et al. 2010), as well as increases in neuronal firing rates of V1 neurons in mice (Shuler and Bear 2006). Stimulation of this region has also been shown to disrupt timing performance, yet only when performed early within the interval (Salvioni et al. 2013), an effect which may rely on individual alpha frequency (Mioni et al. 2020). Curiously, visual cortex activity was only observed on the Initial trials, and not on Re-do, when timing performance improved. One possibility is that, on the Initial trials, subject timing was partially driven by the exogenous visual cue, whereas on the Re-do trials, this cue was more effectively ignored, allowing subjects to rely on an endogenous sensorimotor signal.

For the right hippocampus, this region was also observed primarily on the reproduction phase of Initial trials, which may be due to subjects retrieving the experienced cue from memory. Yet, this region was also less active on the Re-do trials, when subject performance improved. One possibility here may relate to so-called “central tendency” effects commonly observed in time reproduction tasks specifically (Jazayeri and Shadlen 2010) and magnitude-based tasks more generally (Petzschner, Glasauer, and Stephan 2015). In central-tendency, subjects gravitate reproduced responses to the mean of the stimulus-set, an effect suggested to arise from reliance on a Bayesian prior in the face of timing uncertainty. Notably, the central-tendency effect is reduced on Re-do trials (Bader and Wiener 2021), which may be explained by subjects relying on a more precise representation of the perceived interval, and so less on the prior (Cicchini et al. 2012). Relatedly, on a distance reproduction task, similar in nature to this one, the right hippocampus has been shown to correlate with the degree of central tendency (Wiener, Michaelis, and Thompson 2016), suggesting a similar effect may be at play here.

Our functional connectivity analysis also demonstrated a double-dissociation between phases and trial types, with inter-network communication between the DMN and the Timing Network increasing more when estimating a time interval on Initial trials and when reproducing one on Re-do trials. One possible explanation for this difference is that, on Initial trials, subjects may already be aware of their error in their initial response, and so only require network cross-talk again when reproducing the interval on the Re-do trial, when subject performance improved. An implication of this finding is that, if the DMN and Timing Network were to become disconnected, subjects would exhibit no improvements in timing performance on Re-do trials, and moreover would be unable to report how they had erred. Evidence for this dissociation comes from a study of patients with Parkinson’s Disease, who exhibited a difference in precuneus activation on a time reproduction task when tested on or off dopaminergic medication (Dušek et al. 2012).

With regards to specific regions, three nodes that exhibited the broadest connectivity patterns were the superior frontal gyrus and ACC of the DMN, and the cerebellum for the Timing Network. Both the superior frontal gyrus and ACC have been implicated previously in performance monitoring studies. In particular, recent work in both humans and non-human primates have demonstrated that sensorimotor regions, particularly the SMA, lead connections with the ACC for the purpose of monitoring actions for errors (Sarafyazd and Jazayeri 2019; Fu et al. 2022; Bonini et al. 2014). For the cerebellum, a region long implicated in sensorimotor timing, connectivity with the DMN was highest when encoding the stimulus duration. One aspect for this may relate to self-awareness of timing uncertainty as the interval is measured (Peterburs and Desmond 2016). Indeed, recent work has suggested the cerebellum serves to compute a Bayesian posterior estimate of duration (Narain et al. 2018).

## Conclusion

Overall, this study underscores the importance of timing cognizance in improving timing performance. The recruitment of the DMN as the interval duration is encoded and perceived in the Re-do signals this network’s dual responsibility in tracking and adjusting to timing errors while a properly timed motor response is implemented in a situation with multiple opportunities (for example, hitting a baseball on the second try).

Finally, an understanding of how time awareness operates in the neurotypical brain can provide insights when timing is disrupted. In the case of autistic children, time reproduction accuracy and precision remained intact while the children were unable to evaluate their own timing ability (Doenyas et al., 2019). Due to the internetwork connectivity between the default mode and timing networks, any disorder or injury that impacts the TN structures could also potentially adversely affect the DMN regions, leading to not only a distortion in temporal representation but also in its awareness.

